# Managing genomic variant calling workflows with Swift/T

**DOI:** 10.1101/524645

**Authors:** Azza E. Ahmed, Jacob Heldenbrand, Yan Asmann, Faisal M. Fadlelmola, Daniel S. Katz, Katherine Kendig, Matthew C. Kendzior, Tiffany Li, Yingxue Ren, Elliott Rodriguez, Matthew R. Weber, Justin M. Wozniak, Jennie Zermeno, Liudmila S. Mainzer

## Abstract

Genomic variant discovery is frequently performed using the GATK Best Practices variant calling pipeline, a complex workflow with multiple steps, fans/merges, and conditionals. This complexity makes management of the workflow difficult on a computer cluster, especially when running in parallel on large batches of data: hundreds or thousands of samples at a time. Here we describe a wrapper for the GATK-based variant calling workflow using the Swift/T parallel scripting language. Standard built-in features include the flexibility to split by chromosome before variant calling, optionally permitting the analysis to continue when faulty samples are detected, and allowing users to analyze multiple samples in parallel within each cluster node. The use of Swift/T conveys two key advantages: (1) Thanks to the embedded ability of Swift/T to transparently operate in multiple cluster scheduling environments (PBS Torque, SLURM, Cray aprun environment, etc.,) a single workflow is trivially portable across numerous clusters; (2) The leaf functions of Swift/T permit developers to easily swap executables in and out of the workflow, conditional on the analyst’s choice, which makes the workflow easy to maintain. This modular design permits separation of the workflow into multiple stages and the request of resources optimal for each stage of the pipeline. While Swift/T’s implicit data-level parallelism eliminates the need for the developer to code parallel analysis of multiple samples, it does make debugging of the workflow a bit more difficult, as is the case with any implicitly parallel code. With the above features, users have a powerful and portable way to scale up their variant calling analysis to run in many traditional computer cluster architectures.

https://github.com/ncsa/Swift-T-Variant-Calling

http://swift-t-variant-calling.readthedocs.io/en/latest/

## Introduction

Advancements in sequencing technology [1,2] have paved the way for many applications of Whole Genome Sequencing (WGS) and Whole Exome Sequencing (WES) in genomic research and the clinic [3,4]. One of these applications is genomic variant calling, commonly performed in accordance with the Best Practices established by the GATK team (Genome Analysis Toolkit) [5–7]. This methodology involves constructing a complex workflow that could be hard to manage especially for large sample sizes (hundreds and beyond, [8–10]) that necessitate the use of large computer clusters. In such cases, features like resiliency and auto-restart in case of node failures, tracking of individual samples, efficient node utilization, and easy debugging of errors and failures are very important. Without a high-quality workflow manager, these requirements can be difficult to satisfy, resulting in error-prone workflow development, maintenance and execution. An additional challenge is porting the workflow among different computing environments, a common need in collaborative and consortium projects.

Monolithic solutions, where a single executable runs the entire analysis, can replace the complex multi-stage workflow and obviate the need for workflow management. Examples of these solutions include Isaac [11], Genalice [12] and Dragen [13]. These programs offer a plethora of options, but may be too rigid for some analyses, preventing users from swapping algorithms for better accuracy or making adjustments for different species (reference genome, ploidy, known SNP sets etc.) [14]. These monolithic solutions are also developed and maintained by private companies, which may delay or preclude the incorporation of novel approaches and algorithms developed by the scientific and medical community.

It is likely that the GATK will continue to be the standard in research and medicine for those reasons, and also due to the need for HIPAA [15]/CLIA [16] approval and compliance. The GATK is well trusted, validated by the community, and grandfathered in. Thus, the need for a generic, modular and flexible workflow built around the toolkit will persist for some time.

Multiple workflow management systems are now available [17] that differ in their design philosophy and implementation. None so far have been found to be the “best” choice for bioinformatics, although some winners are emerging, such as the Common Workflow Language (CWL [18]) and the Workflow Definition Language (WDL [19]), see Discussion. Key distinguishing features are the underlying language and syntax in which the workflow is expressed, and the monitoring and parallel processing capabilities of workflows while executing. Swift/T [20] is one such workflow management system, composed of Swift - a high-level, general-purpose dataflow scripting language [21], and Turbine - a workflow execution engine [22]. The greatest purported advantages of Swift/T are its high portability and ability to scale up to extreme petascale computation levels [23]. Additionally, a number of features make this language an attractive choice for complex bioinformatics workflows [24]:

- *Abstraction and portability*, where cluster resource management is largely hidden from the user, allowing the same code to be seamlessly ported among clusters with different schedulers;
- *Modularity* through the use of leaf functions to define heavyweight processing tasks that are called as need arises;
- *Extensibility* through easy integration of functions written in other languages;
- *Dataflow-based programming framework* that ensures efficient use of compute resources through compile-time optimization for distributed-memory computing models and hybrid parallelism, resulting in high scalability;
- *Code readability* due to its C-like syntax; and
- *Code expressibility* - inclusion of standard programming features, such as conditional execution, iteration, and recursive functions [25].

We explored Swift/T as a choice in the space of currently available workflow management systems. This paper documents our experience implementing, debugging and deploying a genomic variant calling workflow in Swift/T available at https://github.com/ncsa/Swift-T-Variant-Calling and documented on http://swift-t-variant-calling.readthedocs.io/en/latest/.

## Methods and Results

The GATK Best Practices variant calling workflow consists of multiple steps that require conditional adjustments based on the analysis use case, such as whole genome vs. exome sequencing, paired- or single-end reads, species or ploidy, etc. The primary role of the workflow management system, such as Swift/T, is to handle this conditional branching and coordinate the launch of command-line tools in accordance with the user-defined configuration and data dependencies, while efficiently managing the computational resources. The underlying workflow language should make it easy to develop and maintain such complex workflows. Based on our prior experience in scaling-up the variant calling workflow [26–28], and that of others [29–31], we have put together a list of requirements to be satisfied while redesigning the workflow in Swift/T, and used them to evaluate the performance of the language for our purposes.

## Workflow design requirements

### Modularity

By definition, a workflow is a series of computational tasks, where outputs of one task serve as inputs to the next. Each task can be performed by a selection of bioinformatics software package options driven by the nature of the analysis (Table 1). This flexibility can be enabled by constructing *modular* workflows, such that each executable is incorporated via a generic wrapper, making it easy for the developer to swap executables at the task level. For example, at the level of the *Alignment* task, the workflow language should permit easy swapping of BWA MEM [32] for Novoalign [33], conditionally on an option stated in a configuration or run file.

**Table 1.**
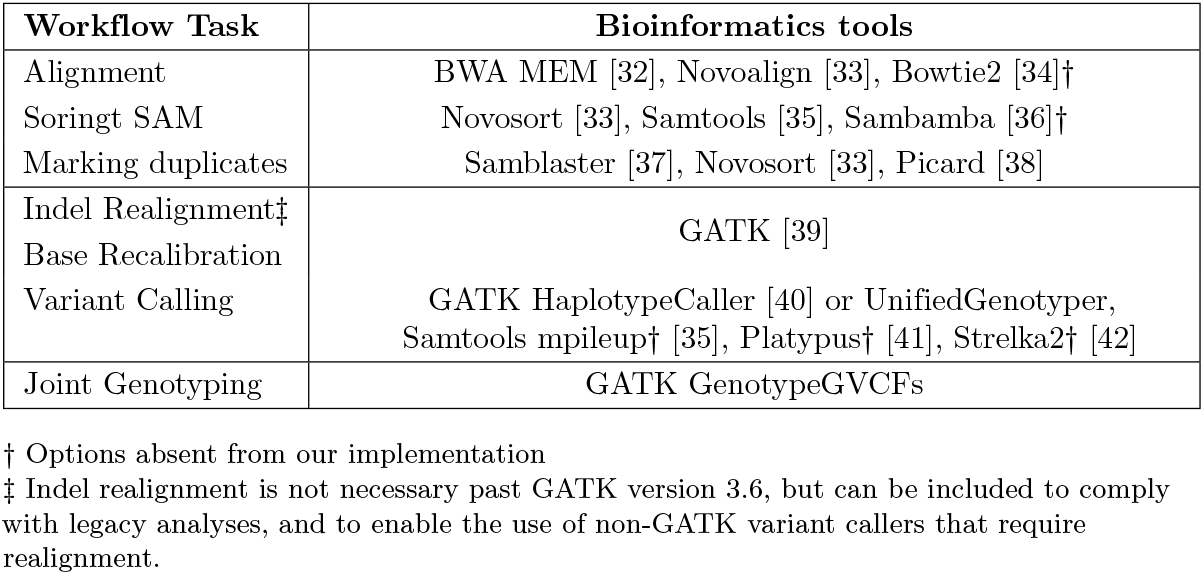
Tools commonly used in genomic variant calling workflows.

| Workflow Task | Bioinformatics tools |
| --- | --- |
| Alignment | BWA MEM [32], Novoalign [33], Bowtie2 [34]† |
| Sorting SAM | Novosort [33], Samtools [35], Sambamba [36]† |
| Marking duplicates | Samblaster [37], Novosort [33], Picard [38] |
| Indel Realignment‡ | GATK [39] |
| Base Recalibration |  |
| Variant Calling | GATK HaplotypeCaller [40] or UnifiedGenotyper, Samtools mpileup† [35], Platypus† [41], Strelka2† [42] |
| Joint Genotyping | GATK GenotypeGVCFs |

† Options absent from our implementation
‡ Indel realignment is not necessary past GATK version 3.6, but can be included to comply with legacy analyses, and to enable the use of non-GATK variant callers that require realignment.

Many tools in Table 1 can take a long time to run on deeply-sequenced samples. This poses a problem for analyses run on computer clusters that have a restrictive maximum job walltime limit. Thus it is useful to break up the workflow into *stages* - integrated sets of tasks that can be viewed as higher-level modules. Each module is then executed as its own cluster job that fits within the maximum walltime constraint. Chaining such modules together into one executable script effectively requires support for “workflows of workflows”.

The modular architecture has additional advantages conferring economy of compute resources and maintainability of code. It allows the user to run a portion of the workflow on the resources optimal for that particular stage, which is useful when a workflow has many fans and merges, but the fans have different node-widths among them. In case of runtime failure, it also enables users to restart the workflow at a failed stage without having to recompute successful upstream calculations. The latter advantage, however, is obviated if the workflow management system itself provides seamless workflow restart from the point of failure - a required feature for complex workflows running at scale. Finally, modularity ensures that the implementation of individual stages can be altered without breaking the workflow, as long as inputs and outputs remain consistent. This way, workflows can be updated with new methodologies as the scientific field and respective tools evolve.

### Data parallelism and scalability

A major expectation of a good workflow management system is the ability to develop a single code path that will automatically run in parallel on multiple samples and not force the user to manually code data-level parallelism. This *implicit* parallelism is not just a matter of convenience, but a significant performance boost. Bioinformatics tools are commonly implemented as multithreaded executables that are not MPI-enabled. Thus, in Bash workflows each task on each sample has to be run as an individual cluster job. If the cluster does not support job arrays, its workload manager can get overwhelmed by the high number of jobs when analyzing large datasets, leading to slow queues or failures. In contrast, a proper workflow management system should run a workflow as a single multi-node job, handle the placement of tasks across the nodes using embedded parallel mechanisms, such as MPI, and scale well with the number of samples.

The workflow manager should also support repetitive fans and merges in the code. For example, in variant calling it is common to cut the walltime of analysis by splitting the input sequencing data into chunks, performing alignment in parallel on all chunks, merging the aligned files per-sample for sorting and deduplication, and finally splitting again for parallel realignment and recalibration per-chromosome (Fig. 1, left panel). This pattern of parallelization is more complex than merely running each task on each input sample - yet is a common workflow requirement.

**Fig 1.**
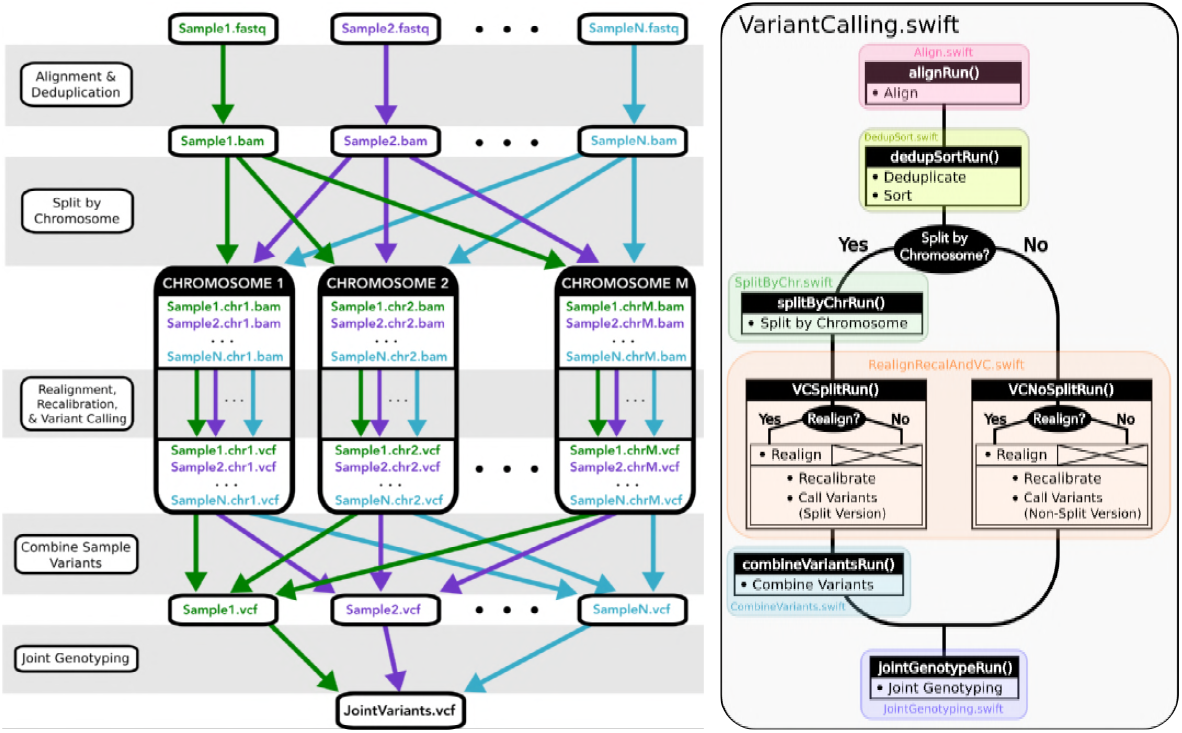
Swift/T variant calling code, under the hood. Left: Patterns of parallelization implemented in our Swift/T variant calling workflow. Right: Colored blocks represent the different stages of the workflow. Black blocks indicate methods within the respective modules.

Finally, in bioinformatics we only need certain tools to run on multiple samples in parallel. Other tasks, such as creating folders, user notification or running QC on the whole stage, can and sometimes should be run sequentially. Therefore, it is beneficial to support differential use of data-level parallelism in some modules but not others.

### Real-time logging and monitoring

When analyzing many samples at once, especially in a production environment where the data flow continuously through the cluster, it is important to have a good system for logging and monitoring progress of the jobs. At any moment during the run, the analyst should be able to assess (1) which stage of the workflow is running for every sample batch, (2) which samples may have failed and why, (3) which nodes are being used by the analysis, and their health status. Additionally, a well-structured post-analysis record of all events executed on each sample is necessary to ensure reproducibility of the analysis. This can be manually accomplished by developing a system of runtime logs captured via `stdout` dumps, and handling user notification via `mailx`, but both are quite tedious to code for complex, branched, multi-task workflows. A good workflow manager should provide these capabilities implicitly.

### Portability

A developer should be able to write a workflow once and then deploy it in many environments: clusters with different node configuration, multiple queues and job schedulers, in HPC or in the cloud. For a workflow as complex as genomic variant calling, having to change and adapt for each different cluster is extremely counterproductive.

## Implementation of Design Requirements in Swift/T

### Modularity

The Swift/T language natively supports modularity by defining a “worker” for each executable (“leaf function” in Swift/T terminology), to be called at the appropriate place in the workflow. For example, we implemented the choice to align reads either using BWA MEM or Novoalign, as follows.

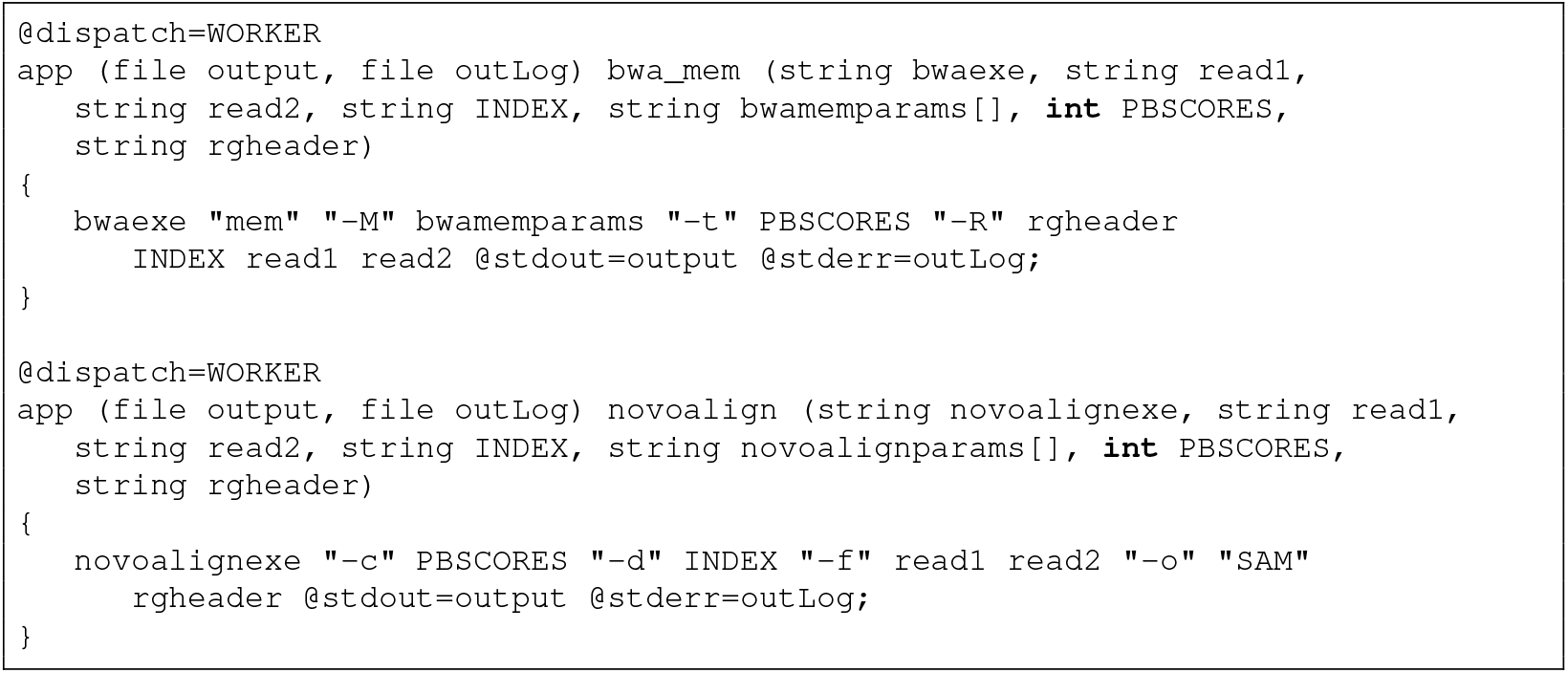

Here each executable is wrapped using the generic “worker” syntax, and workers are conditionally invoked in a compact fashion to perform the Alignment task of the workflow.

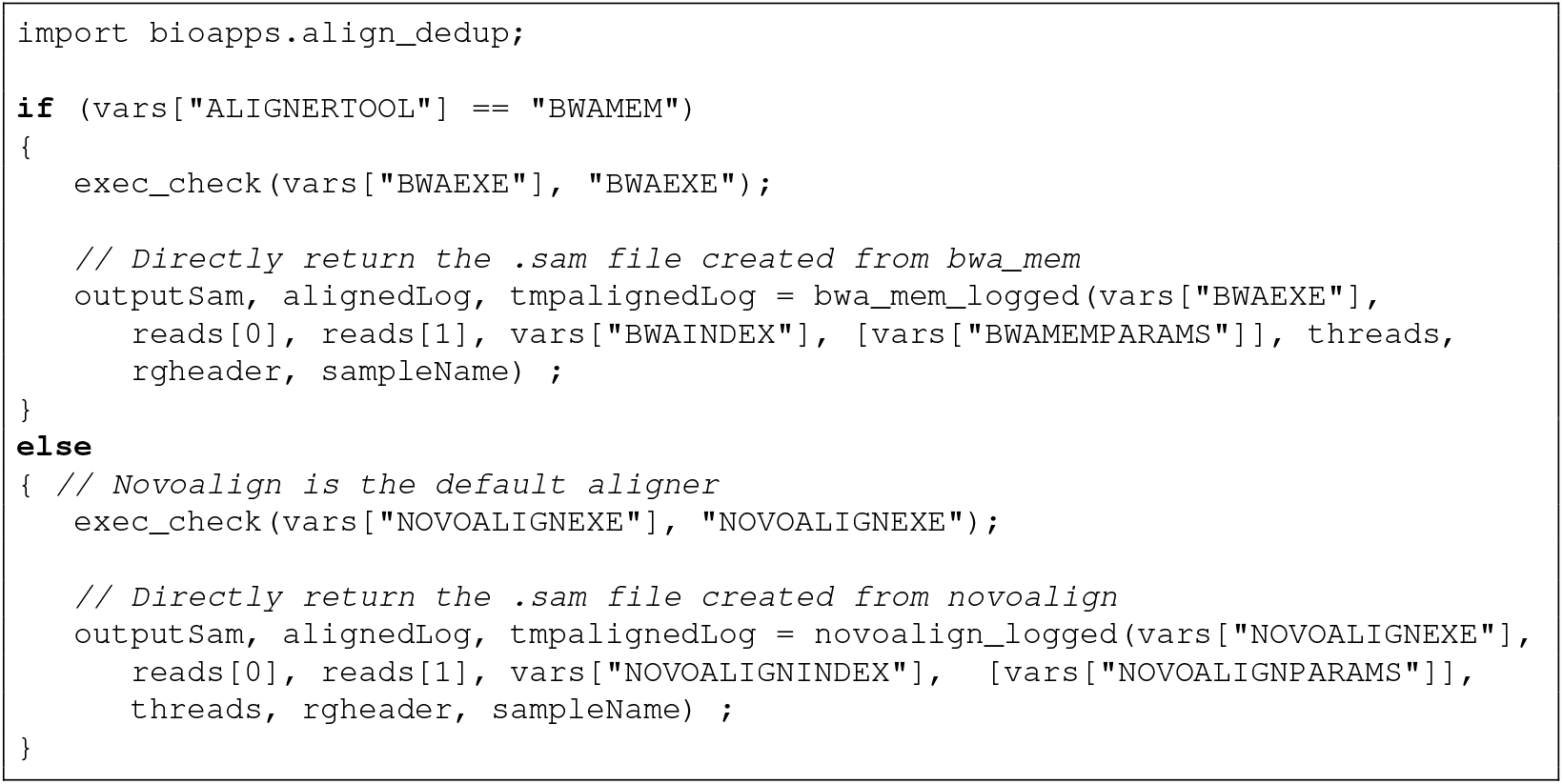

Subworkflows, or “stages”, are implemented as individual Swift/T app functions that are chained together by the primary workflow script (Fig 1, right panel). At each stage, the user can direct the workflow to generate the output files necessary for the next stage, or pass on the output generated from a previous run. At the end of each stage, there is an implicit wait instruction that ensures all tasks have finished before the next stage can run (also see next section).

### Data parallelism and scalability

The “data flow” programming model of Swift/T implicitly supports parallel execution of tasks. Statements are evaluated in parallel unless prohibited by a data dependency or resource constraints, without the developer needing to explicitly code parallelism or synchronization. Swift/T will automatically wait on a process to finish if the next step depends on its output. For example, after read alignment, the step to mark duplicates in an aligned BAM (`picard_logged`) depends on the previous step (`novosort_logged`), which produces a sorted BAM (`alignedsortedbam`) to serve as input to the deduplication step. The essense of implicit parallelization is that `picard-logged` will wait until `novosort_logged` is finished due to this data dependency.

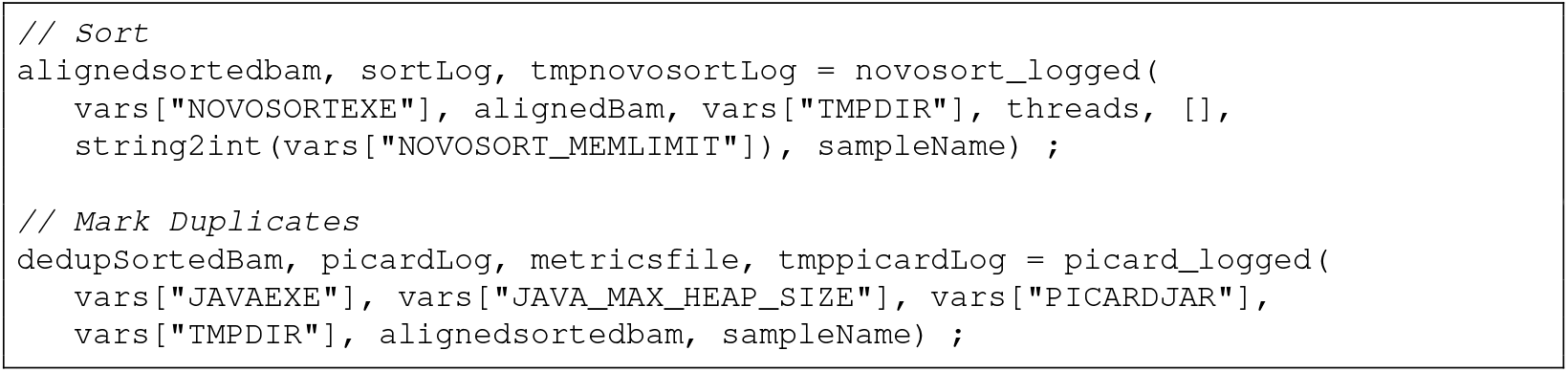

There are some places in the workflow where a stage must wait on another, yet a direct data dependency does not exist. For example, log information begins to be produced right away as the Alignment module begins execution. The output log folder must first exist for this purpose, but the asynchronous parallel execution function of Swift/T may start the Alignment module before it runs the statement to create the log folder. This can be addressed by explicitly forcing the wait either via the “=>“ symbol, `via wait ()` statement, or via a dummy variable that “fakes” a data dependency.

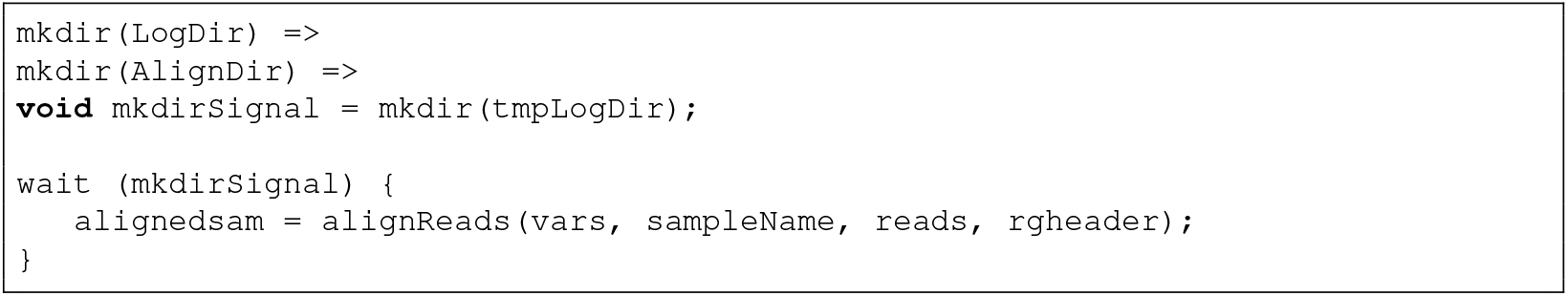

The above example illustrates the use of a `wait()` statement, and also the drawbacks of enforcing implicit parallelism across the entire workflow. In bioinformatics, patterns of execution are usually mixed: individual commands running in parallel on many samples are intermixed with serial blocks of code that perform quality control, data management, user notification, or other tasks. It would be useful to have these blocks fenced-off to prevent Swift/T from attempting to run them all asynchronously and in parallel. Parsl, the next step in evolution of Swift language, has that capability [43,44].

Nonetheless, Swift/T does take care of parallelism in a smart and transparent way that makes efficient use of resources. The user should still take care to request a reasonable number of nodes: too few - and many samples will be processed in series; too many - and resources will be reserved unnecessarily. Beyond that there is no need to worry about task placement, as Turbine will take care of it. This is extremely useful, because bioinformatics programs do not always scale well to the full number of cores available on the compute nodes, and therefore running multiple instances of a task simultaneously on the same node may improve the overall efficiency. For example, BWA MEM normally scales well up to eight threads, so running two eight-thread processes in parallel on a 16-core node is more efficient than running two sixteen-thread processes in series. We implemented this as user-level options that specify the number of cores per node and the number of programs to run on each node simultaneously. From there the workflow determines the number of threads to use for each bioinformatics program, and Swift/T uses Asynchronous Dynamic Load Balancing (ADLB) [45] to distribute those programs across nodes as they become available at run time. Without ADLB one would have to code this explicitly for each job scheduler, which becomes very complicated on clusters that do not support node sharing, i.e. only one job is allowed to run per node. In the latter case a vanilla Bash workflow [46] would need to incorporate an MPI wrapper (e.g. [47]) to take care of program placement across nodes. The MPI backend of ADLB fulfills that function in Swift/T.

We verified correctness of the task dependency chains and parallel execution by tracking start and end times of each task for multiple samples in some of our tests (see next section and Fig. 2).

**Fig 2.**
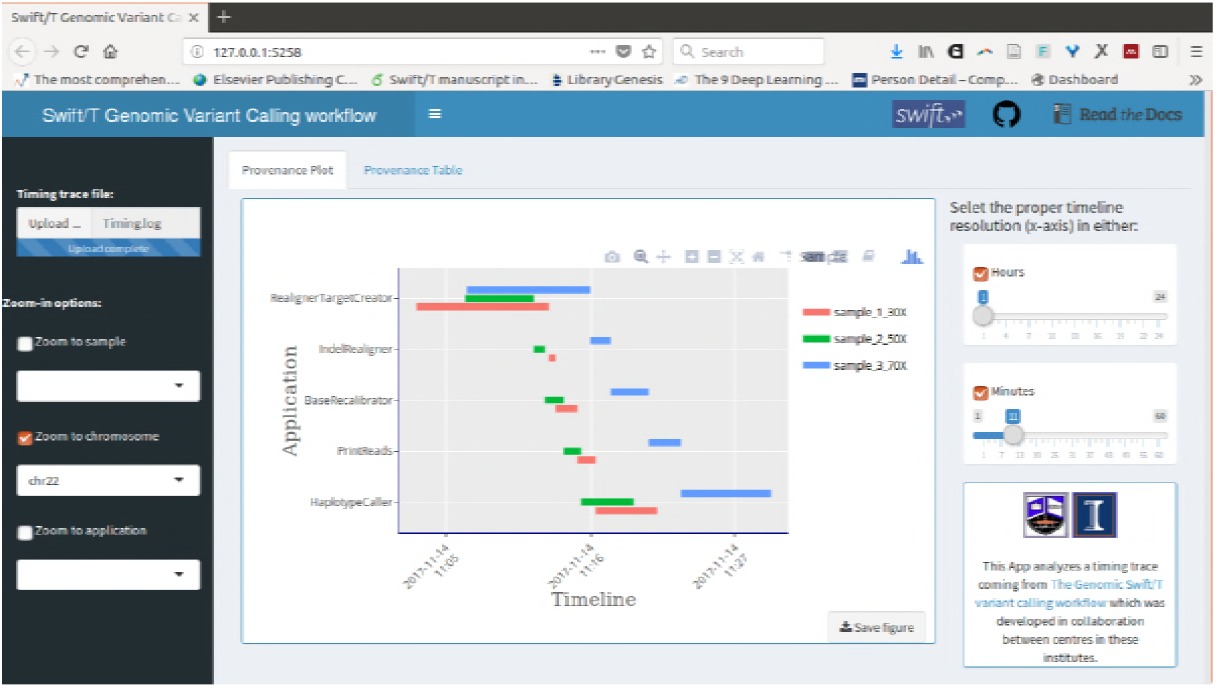
Timing provenance tracking of a 3-sample pipeline run (synthetic whole exome sequencing dataset at 30X, 50X and 70X) on Biocluster [50]. This plot view is interactive, allowing full pan and zoom and was generated using plotly library in R.

### Real-time Logging and Monitoring

The underlying MPI-based implementation of Swift/T logic makes it possible to leverage standard MPI logging libraries to collect run-time details about the status of every sample. We used the Message Passing Environment (MPE) library [45] to log the usage of the MPI library itself and ADLB calls [48], and implemented visualization in Jumpshot viewer. To enable such logging requires installation of the MPE library in addition to the standard Swift/T components (C-utils, ADLB library, Turbine and STC). This turned out to be a bit cumbersome because it requires creation of new functions: tcl wrappers around MPE to log when each executable starts and stops.

Another approach to tracking the workflow run time execution is to manually implement Swift/T leaf functions such that the start and end timing of each function are logged. A timing graph can be generated using R script based on this information, showing the analysis steps across samples, chromosomes and specific applications (Fig 2). Interactivity is added via Shiny R package [49]. This is a fairly manual approach, little better than the Bash echo date statements. Nonetheless, it permits one to view the patterns of pipeline execution even if it fails, and partial logs can similarly be viewed as the pipeline is running. To obtain the up-to-date trace, one can type in the R terminal:

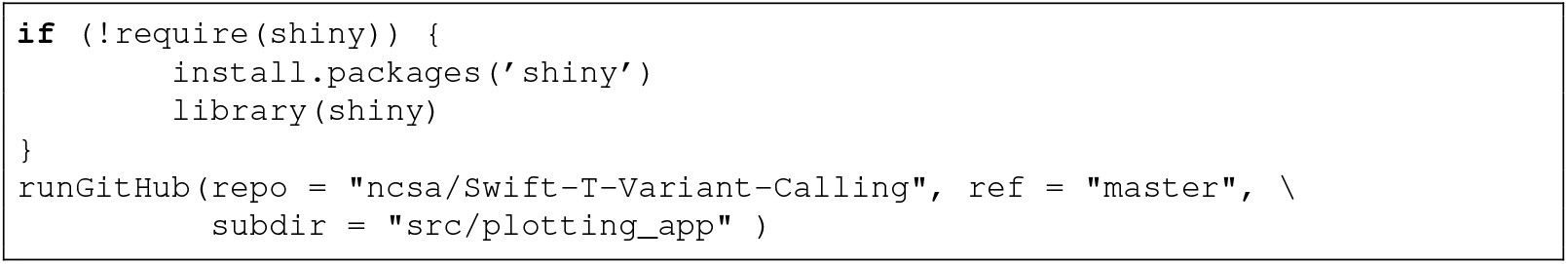

In conclusion, logging and monitoring can be usefully implemented in a Swift/T workflow, but are not adequately supported at the time of this writing and require quite a bit of work.

### Portability

Swift/T runs as an MPI program that uses the Turbine [22] and ADLB [45] libraries to manage and distribute the workflow execution on local compute resources (desktop/laptop), parallel computers (clusters/HPCs), and distributed systems (grid/cloud). Its built-in wrappers can launch jobs on many common resource schedulers, such as PBS Torque, Cobalt, Cray aprun, and SLURM [51], using the `−m` flag passed to the Swift/T executable, i.e. `swift−t −m slurm`. Through these unified wrappers, the user is only left with the trivial task of specifying the required computational resources: queue, memory, wall time, etc.:

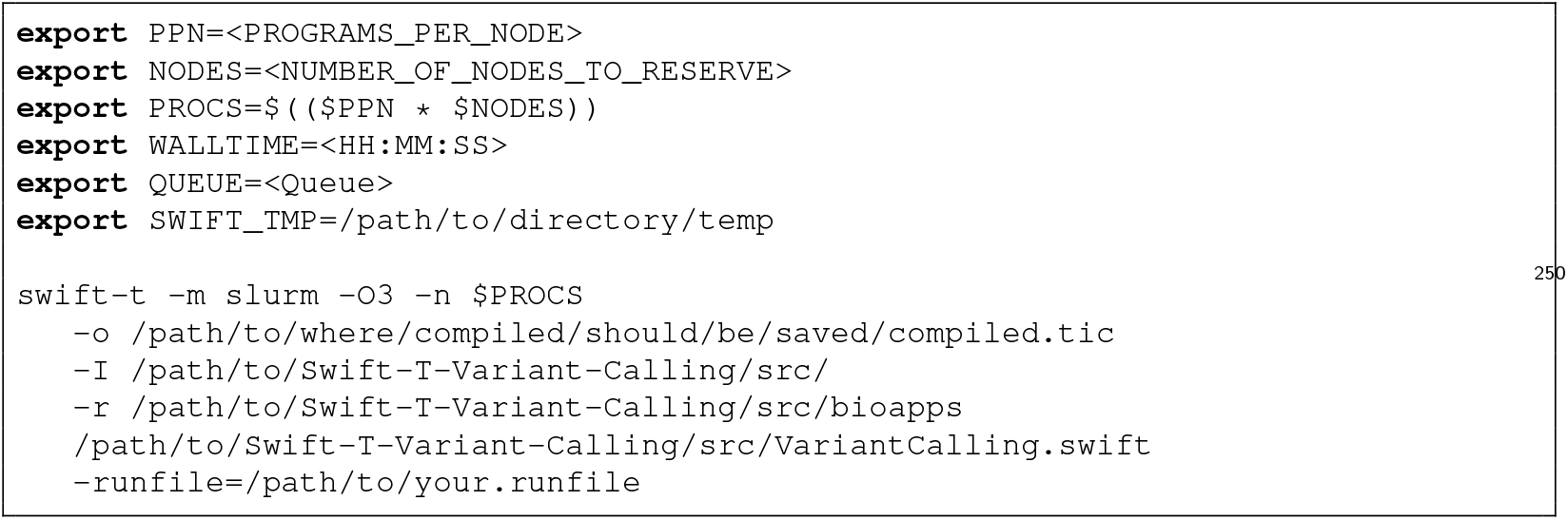

We verified both portability and scalability conferred by Swift/T by testing on a variety of HPC systems with a range of cluster setups, job schedulers and patterns of execution (Table 2). Portability across resource schedulers works as expected, although unique setups may require tweaks, such as setting of environmental variables [52], with configuration of 1 sample/node and 2 samples/node.

**Table 2.** Swift/T delivers on its promise of portability and scalability. Synthetic data were generated using the NEAT synthetic read simulator [54]. Node sharing column indicates whether the cluster permits jobs to share the same node.

| System | Resource manager | Node type | # nodes per run | Node sharing | Test data |
| --- | --- | --- | --- | --- | --- |
| iForge [55] | PBS Torque | IvyBridge, 20 cores, 256 GB RAM | 1-8 | No | Soy NAM [56] using 2, 6, 12, or 16 sample batches † |
| XSEDE Stampede2 [57] | Slurm | KNL, 68 cores, 4 hardware threads/core, 96 GB DDR4, 16 GB MCDRAM | 1 | Yes | GIAB NA12878 Illumina HiSeq Exome (NIST7035) [58]; Synthetic chr1 exome seq 50X |
| Biocluster [50] | Slurm | Dell PowerEdge R620, 24 Cores, 384 GB RAM | 1; 3 | Yes | Synthetic WES 30X; Synthetic WES 50X; Synthetic WES 70X |
| Single server at CBSB, H3ABioNet node | N/A | HP Proliant dl380p gen. 8 24 cores 125 G RAM | 1 | Yes | Synthetic chr1 exome seq 50X |

† This Swift/T variant calling workflow was also used on iForge for a variety of analyses on WES and WGS data in different species.

All other functionality of our workflow was also fully validated on soybean and human Illumina sequencing data, as well as synthetic datasets. The complete list of tested options and features can be found on our GitHub repository [53].

### Robustness against failure

Swift/T has native support for restarting a task after failure. The user controls the maximum number of allowed retries, and a randomized exponential backoff delay is applied between them, attempting to rerun the task until success or the pipeline terminates, whichever is sooner. Retries do not correct for bugs in the pipeline code, but only for Swift/T leaf function failures that are not related to compilation errors or “assert” failures.

This is useful when applications fail for nondeterministic reasons, such as a filesystem under load slowing down I/O and making the application wait for data, thus causing it to time out. However, when running wide jobs on large clusters, it is also necessary to have robustness against node failure. In collaboration with the Swift/T team, we introduced the support for moving the retries of the failed task to another, randomly chosen, MPI rank. For reproducibility purposes, random number generation in Swift/T defaults to start from the same seed, which is dependent on the MPI rank where the process is to be evaluated, unless the seed is specified by the turbine variable “`TURBINE_SRAND`”.

## Discussion

Complexity of problems in biology means that nearly every kind of analytics is a multi-step process, a pipeline of individual analyses that feed their outputs to each other (e.g. [59–61]). The algorithms and methods used for those processing steps are in continual development by scientists, as computational biology and specifically bioinformatics are still rapidly developing. Few studies can be accomplished via a single integrated executable. Instead we deal with a heterogeneous medley of software of varied robustness and accuracy, frequently with multiple packages available to perform seemingly the same kind of analysis - yet subtly differing in applicability depending on the species or input data type. Thus bioinformatics today requires advanced, flexible automation via modular data-driven workflows. This is a tall order, considering the added requirements of scalability, portability and robustness. Genomics is a big data field: we no longer talk about sequencing individual organisms, but every baby being born (~500 per day per state in the US) and every patient who comes in for a checkup (a million per year in a major hospital), not to mention the massive contemporary crop and livestock genotyping efforts. The workflows managing data analysis at that scale must take full advantage of parallelism on modern hardware, be portable among multiple HPC systems and the cloud, be robust against data corruption and hardware failure, and provide full logging and reporting to the analyst for monitoring and reproducibility.

Recently there has been an incredible upsurge in developing scientific workflow management systems, enough to have resulted in calls for standardization and quality assurance [62]. In this manuscript we reviewed our experience with one such system, Swift/T, touching on workflow management, performance and scalability issues; security was deemed out of scope.

### Pros and cons of Swift/T for bioinformatics workflows

Swift/T is a powerful and versatile language that offers many advantages for production large-scale bioinformatics workflows. It allowed us to fulfill most of the requirements outlined in the *Requirements* section, for variant calling workflow as a use case. Below is our summary of pros and cons based on that experience.

*Portability* may well be the greatest strength of Swift/T: a workflow written in Swift/T can be executed on a wide variety of compute infrastructures without changing the code, and the user does not need to know about the underlying scheduling environment on the cluster. The language abstracts away the low level concerns such as load balancing, inter-process communication and synchronization of tasks automatically through its compiler (stc) and runtime engine (Turbine), allowing the programmer to focus on the workflow design [63].

*Modularity* is another excellent advantage of Swift/T. The language glues together command line tools: either directly by wrapping them in Swift/T app functions if they solely operate on files; or indirectly as tcl packages with corresponding Swift/T app function declarations if they produce numerical or string outputs. Under the hood, Swift/T code is actually compiled into Tcl syntax before Turbine gets to manage the distribution and execution of tasks to compute resources. This further means that wrapping any C, C++ or Fortran application is also easy due to Tcl. This leaf-function modularization, and the ease of integrating code written in other languages into Swift/T environment, is the reason why we preferred this to its predecessor Swift/K [21], which had superior provenance and checkpointing capabilities [64].

*Implicit data parallelism and scalability* of Swift/T is a powerful way of enabling big data analyses by increasing the amount of simultaneous computation. The language particularly lends itself to use cases that require asynchronous rapid-fire of small, quick parallel jobs [65]. That is one of the many kinds of bioinformatics workloads, but not the most typical one for primary analysis of genomic data. In this field we frequently require a simple wrapper to run a single, time-consuming step on a large number of samples or other units of data level parallelization: i.e. conversion of several thousand BAMs back to FASTQs for reanalysis with the most recent reference genome. However, the data flow task parallelism framework has a substantial learning curve, despite offering familiar control flow statements and expressions in C-like syntax [66]. Coding and debugging can require a more substantial effort than say, Nextflow [67], and that can be a barrier for biologists. An additional inconvenience is that Swift/T does not support piping between applications, which is extensively used in bioinformatics analyses, as they are still overwhelmingly file-based pipelines.

*Robustness against failures* in Swift/T is supported via leaf function retries, attempting to rerun the task on one of the available ranks. This confers resilience against nondeterministic failures, such as filesystem or cluster interconnect hiccups as well as hardware failures - an important advantage for big data genomics.

*Real time logging* is provided via runtime Turbine logs, with user-controlled verbosity. These can be quite detailed but challenging to use for debugging when the analyst must understand whether a failure occurred due to data, a bioinformatics application or the Swift/T code bug. The greatest difficulty stems from asynchronous log records, caused by asynchronous execution of tasks. Thus an error printout rarely corresponds to the execution message that immediately precedes it in the log, and finding the failed tasks from the log alone is nearly impossible. We had to manually implement the per-task and per-executable logs in our code, to counteract this inconvenience.

In summary, Swift/T language lends itself to creating highly portable, modular and implicitly parallel workflows. It is very powerful, especially when a workflow consists of raw code pieces written in C, C++, Fortran, etc. However, it may be overkill for those bioinformatics workflows that consist of pre-compiled executables glued together. The lack of support for piping between applications is a major drawback for big-data bioinformatics, resulting in proliferation of intermediary files. Portability, the main advantage of Swift/T, could perhaps be accomplished in simpler ways. In the following sections we review other workflow management systems, to put Swift/T into the broader context of life sciences.

### Challenges in building the “right” workflow manager for computational biology

The implementation of workflow management systems (WMS) for computational biology, bioinformatics and genomics is strongly influenced by culture and prevailing expertise in the multidisciplinary fields. One has to contend with two populations of scientists: those with strong biology background, driven to solve research problems, to whom programming is an unavoidable yet joyless burden; and those able to produce complex and capable code that is not perhaps very user-friendly. This creates a real problem with adoption of any software, including a WMS: the harder it is for a scientist to use a software package compared to an ad-hoc hack, the lower its widespread acceptance in the community [62]. Perhaps that’s why simple glue solutions via Bash, Perl, Python, Make, CMake and similar, have persisted for so long. Their shallow learning curve permits quick production of short-term analytic solutions, which get used over and over despite poor scaling with growing dataset size, and despite requiring a lot of work to port among compute systems.

Scientific Workflow Systems are the next step up from scripting. Those that provide a graphical user interface, such as Taverna [68], Galaxy [69] and Kepler [70] (Table 3), have good accessibility for scientists with less programming experience but require quite a bit of effort to be set up and maintained, and have limited set of features. In contrast, lower level systems with a command-line interface (CLI), such as Snakemake [71], Luigi [72], BcBio [73], Bpipe [74], are easier to maintain and share, provide good documentation and reproducibility, fault tolerance, and task automation; however, they require a lot more programming expertise.

**Table 3.**
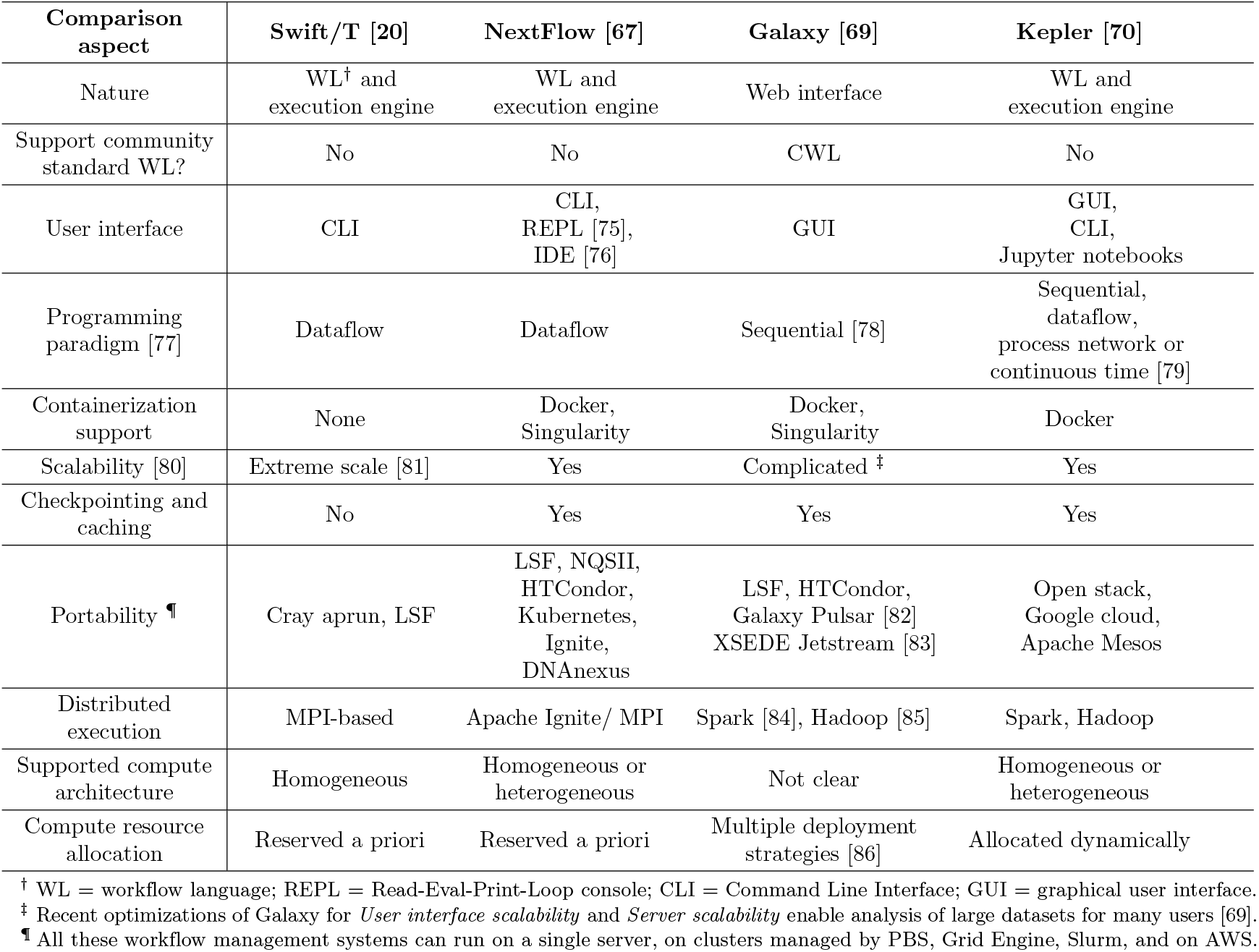
Popular workflow management systems.

| Comparison aspect | Swift/T [20] | NextFlow [67] | Galaxy [69] | Kepler [70] |
| --- | --- | --- | --- | --- |
| Nature | WL<sup>†</sup> and execution engine | WL and execution engine | Web interface | WL and execution engine |
| Support community standard WL? | No | No | CWL | No |
| User interface | CLI | CLI, REPL [75], IDE [76] | GUI | GUI, CLI, Jupyter notebooks |
| Programming paradigm [77] | Dataflow | Dataflow | Sequential [78] | Sequential, dataflow, process network or continuous time [79] |
| Containerization support | None | Docker, Singularity | Docker, Singularity | Docker |
| Scalability [80] | Extreme scale [81] | Yes | Complicated <sup>‡</sup> | Yes |
| Checkpointing and caching | No | Yes | Yes | Yes |
| Portability <sup>¶</sup> | Cray aprun, LSF | LSF, NQSII, HTCondor, Kubernetes, Ignite, DNAnexus | LSF, HTCondor, Galaxy Pulsar [82] XSEDE Jetstream [83] | Open stack, Google cloud, Apache Mesos |
| Distributed execution | MPI-based | Apache Ignite/ MPI | Spark [84], Hadoop [85] | Spark, Hadoop |
| Supported compute architecture | Homogeneous | Homogeneous or heterogeneous | Not clear | Homogeneous or heterogeneous |
| Compute resource allocation | Reserved a priori | Reserved a priori | Multiple deployment strategies [86] | Allocated dynamically |

<sup>†</sup> WL = workflow language; REPL = Read-Eval-Print-Loop console; CLI = Command Line Interface; GUI = graphical user interface.
<sup>‡</sup> Recent optimizations of Galaxy for *User interface scalability* and *Server scalability* enable analysis of large datasets for many users [69].
<sup>¶</sup> All these workflow management systems can run on a single server, on clusters managed by PBS, Grid Engine, Slurm, and on AWS.

The cultural gap in capabilities between developers and end users can be closed via implementation of visual programming (GUI-like interface with CLI-like capabilities), thus allowing for customization of analytic tools and technologies with little to no programming background. But, ultimately the right approach to providing scalability and interoperability is probably via implementation of generic low level bioinformatics specific libraries to be used universally across different sets of tools [87].

In the meantime, great strides are being made by the community in trying out different approaches to scientific workflow management and automation, aiming to satisfy the complex requirements [17]:

- seamlessly managing both serial and parallel steps without creating data waits and computational bottlenecks;
- managing complex task dependencies via explicit configuration (e.g. a user-produced XML file in Pegasus [88]), language-specific syntax (BigDataScript [89]), automatic construction of workflow graphs (Swift [21], WDL [19], Nextflow [67]), rule-based approaches (Ruffus [90] and bpipe [74]) or implicit conventions, while abstracting away from HPC cluster management concerns (Job Management System [91]);
- flexibility to work with varied software being run by the workflow (i.e. via containerization), and widely variegated parameter values and configurations (i.e. through workflow autogeneration [92]);
- ability to handle both fixed and user-defined parameters.

The field seems to have converged on a set of relatively widely used workflow languages (WL) to describe the actual flow of computation, and execution engines (EE) that provide automation and portability on HPC environments. Some solutions are by their nature an integrated package of WL+EE (Table 3). However, there has been a widespread recognition of the need to standardize WLs, for the sake of reproducibility - particularly important for clinical applications. Thus separating out an execution engine that could operate on workflows written in a variety of WLs is very attractive. A few clear leaders have recently emerged: CWL [18] and WDL [19] for workflow definition languages, and Toil [93,94], Rabix [95] and Cromwell [19] for execution engines (Table 4). CWL enjoys very wide adoption, either being supported, or upcoming support announced among Taverna [68], Galaxy [69], Toil [93], Arvados [96], Rabix [95], Cromwell [19]. To some extent such data-driven workflow languages as CWL and WDL can be viewed as a more advanced step in evolution of a formal scientific workflow. Indeed, when a scientist is only experimenting with the new analysis, it is useful to program it in a powerful lower-level language like Swift, which allows a lot of experimentation with the structure and content of the workflow. Once this has been developed and validated, formalizing it in more rigid data-driven framework (CWL, WDL) for reproducibility and later use by non-programmers has a lot of value.

**Table 4.**
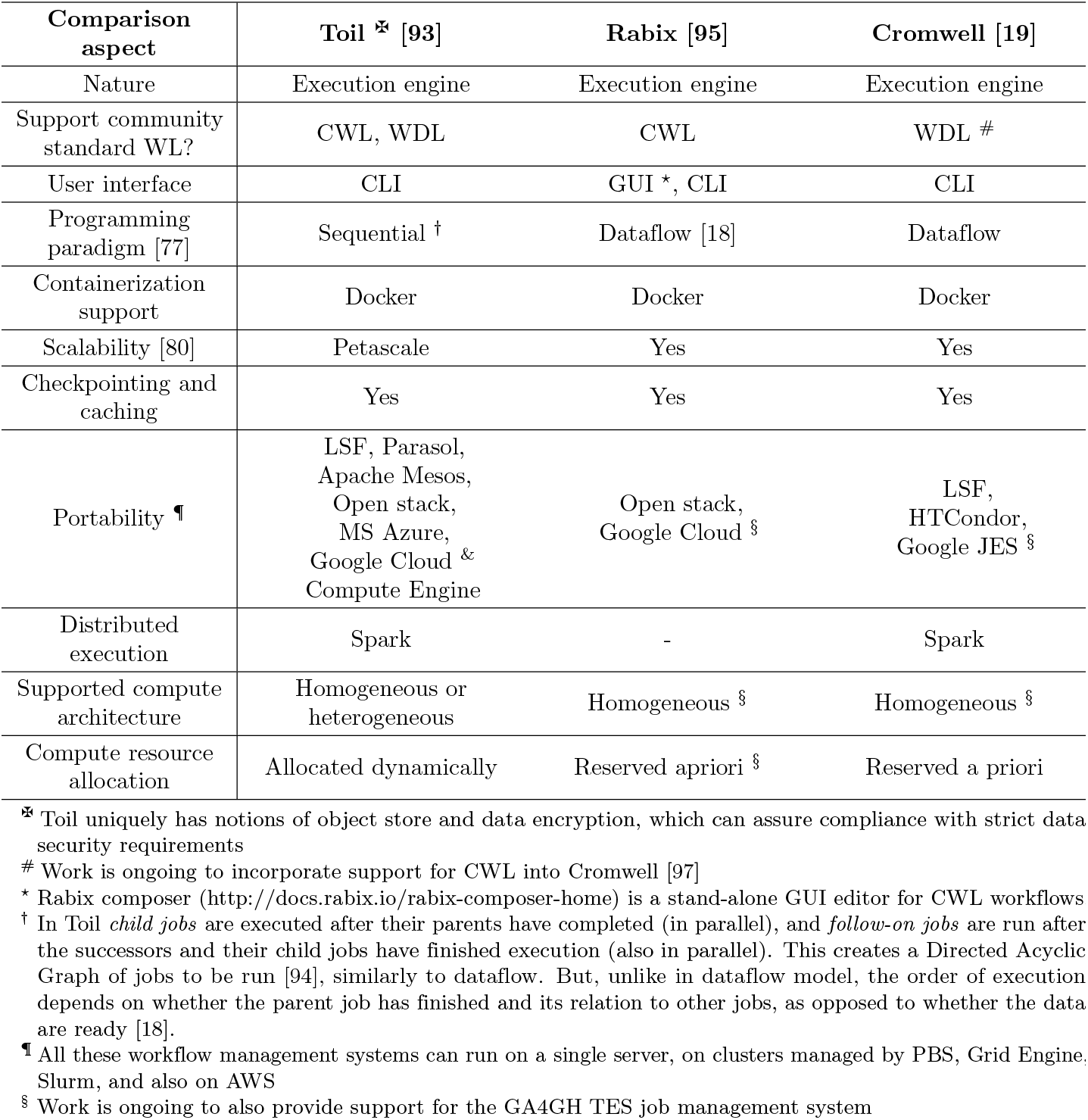
Popular workflow management systems.

Further efforts toward wider adoption recognize the need to execute biomedical workflows on big data platforms, such as Hadoop and Spark (e.g. Luigi), and the cloud (e.g. Toil, DNAnexus, SevenBridges, Illumina’s BaseSpace, Curoverse’s Arvados and iPlant Collaborative’s Agave).

## Conclusion

Our experience implementing a genomic variant calling workflow in Swift/T suggests that it is a very powerful system for workflow management in supercomputing environments. The language is rich with features that give developers control over their workflow structure and execution while providing familiar syntax. The execution engine also has intelligent mechanisms for task placement and regulation, permitting efficient use of compute resources. This unfortunately comes at the cost of a relatively steep learning curve - a common trade-off for programming languages in general. Thus Swift/T can be an extremely useful - and possibly the best - tool for certain genomics analyses, though its complexity may pose an adoption barrier for biologists.

## Acknowledgments

We are grateful for the support of the Blue Waters team, NCSA Industry, and the Argonne/U. Chicago Swift/T developer team during the implementation, testing, and scalability efforts in this project.

This research is part of the Blue Waters sustained-petascale computing project, which is supported by the National Science Foundation (awards 0CI-0725070 and ACI-1238993) and the state of Illinois. Blue Waters is a joint effort of the University of Illinois at Urbana-Champaign and its National Center for Supercomputing Applications.

This work used the Extreme Science and Engineering Discovery Environment (XSEDE), which is supported by National Science Foundation grant number ACI-1548562.

This work used Biocluster, the High Performance Computing (HPC) resource for the Carl R Woese Institute for Genomic Biology (IGB) at the University of Illinois at Urbana-Champaign (UIUC). We are grateful for the support by the Computer Network Resource Group (CNRG) while testing the pipeline.

AEA and FMF are H3ABioNet members and supported by the National Institutes of Health Common Fund under grant number U41HG006941. The content is solely the responsibility of the authors and does not necessarily represent the official views of the National Institutes of Health.

